# Precocious hippocampal structural and functional changes in gestational protein-restricted male elderly offspring: an Alzheimer-simile disease model?

**DOI:** 10.1101/2021.04.07.438796

**Authors:** Gabriel Boer Grigoletti-Lima, Marcelo Gustavo Lopes, Ana Tereza Barufi Franco, Aparecida Marcela Damico, Patrícia Aline Boer, José Antonio Rocha Gontijo

**Affiliations:** Fetal Programming and Hydroelectrolyte Metabolism Laboratory, Medicine and Experimental Surgery Center, Internal Medicine Department, Faculty of Medical Sciences at State University of Campinas, Campinas, SP, Brazil

**Keywords:** Fetal Programming, Alzheimer’s Disease, Behavior and memory, Maternal protein restriction, Hippocampus, β-amyloid peptide, Tau protein

## Abstract

**Background:** Maternal undernutrition has been associated with psychiatric and neurological disorders characterized by learning and memory impairment. Considering the lack of evidence for this, we aimed to analyze the effects of gestational protein restriction on learning and memory function later in life. This research associates behavioral findings with hippocampal cell numbers and protein content related to neurodegenerative brain disease.

**Methods:** Experiments were conducted in animals subjected to a low-protein (LP, 6% casein) or regular-protein (NP, 17% casein) diet throughout their pregnancy. Behavioral tests, isolated hippocampal isotropic fractionator cell studies, immunoblotting, and survival lifetime tests were performed. The results confirmed that the birthweight of LP male pups significantly reduced relative to NP male pups and that hippocampal mass increased in 88-week-old LP compared to age-matched NP offspring. We used the Morris water maze proximity measure, which is the sum of 10 distances each second between rat position and location of a hidden platform target, as a suitable test for assessing age-related learning or memory impairment in aged offspring.

**Results:** The results showed an increased proximity measure in 87-week-old LP rats (52.6 × 10^4^ ± 10.3 × 10^4^ mm) as compared to NP rats (47.0 × 10^4^ ± 10.6 × 10^3^ mm, p = 0.0007). In addition, LP rats exhibited anxiety-like behaviors compared to NP rats at 48 and 86 weeks of life.

Additionally, the estimated neuron number was unaltered in LP rats; however, glial and other cell numbers increased in LP compared to NP rats. Here, we showed unprecedented hippocampal deposition of brain-derived neurotrophic factor, β-amyloid peptide (Aβ), and tau protein in 88-week-old LP compared to age-matched NP offspring. To date, no predicted studies showed changes in hippocampal neuron and glial cell numbers in maternal protein-restricted elderly offspring. The current data suggest that maternal protein restriction has a high impact on lifespan and brain structure, and function.

**Conclusion:** the gestational protein restriction may accelerate hippocampal function loss, impacting learning/memory performance, and supposedly developing diseases similar to Alzheimer’s disease (AD) in elderly offspring. Thus, we propose that maternal protein restriction could be a probable, elegant, and novel method for constructing an AD-like model in adult male offspring.

## INTRODUCTION

In humans and animal models, increasing evidence supports the hypothesis that disturbances during critical periods of fetal development may determine structural and functional alterations later in organs and systems, thereby predisposing individuals to metabolic, cardiovascular, and psychiatric diseases in adulthood [1–6]. As shown in several experimental models, gestational psychological, and nutritional stresses are involved in fetal programming [7–12].

In the embryonic phase, both cell proliferation and the differentiation of vital organs and neural connections are successfully achieved with maternal nutrition. Many neuroanatomical studies investigating the effects of prenatal malnutrition have found decreased neuronal body size, loss of neurons, reduced apical dendrites, branching, and reduced spine density in the hippocampus CA3 pyramidal cell layer [13–16]. Besides, considerable losses of neurons and synaptic ends [17] lead to progressive and, consequently, functional memory loss.

The hippocampus is a brain structure known to be related to emotional behavior; it is crucial for specialized learning and the process of forming memories. In addition, although hippocampal synaptic plasticity is involved in these functions, the details of this slight modulation are not entirely understood [18]. Alzheimer’s Disease (AD) presents specific pathological brain hallmarks represented by extracellular deposition of the β-amyloid peptide (Aβ) aggregates in senile plaques and neurofibrillary tangles (NFT) from excessive tau phosphorylation in different regions of the brain, including the hippocampus.

In human studies, during World War II (1944–1945), the Dutch famine showed a possible relationship between persons exposed to famine during the second trimester and affective psychosis, lending plausibility to reports associating affective psychoses with prenatal exposure [19]. As another critical moment of recent human exposure during gestation to undernutrition, the Chinese famine period, lasting more than three years (1959-1961), promoted a significant incidence of lower cognitive performance and brain development and mental disorders in adulthood [20]. Thus, a low intrauterine nutritional environment has been associated with severe psychiatric and neurological disorders characterized by learning and memory dysfunction, schizophrenia, autism, and neurodegenerative diseases [21].

These relevant neuropsychological disorders have led researchers to develop genetically modified animals that could rescue the disease’s main characteristics and non-genetically modified animals, complementing these models. Considering the lack of evidence, we aimed to analyze the effects of gestational protein restriction (LP) offspring compared to regular protein intake age-matched progeny (NP) on learning and memory functions later in life. In addition to behavioral tests, isotropic fractionator study was performed to evaluate the hippocampal cell number and immunoblotting to determine the content of AD development-related proteins in gestational protein-restricted elderly male offspring.

## MATERIALS AND METHODOLOGY

### 1. Animals

Experiments were conducted on age-matched sibling-mated Wistar HanUnib rats (250–300 g). General guidelines established by the Brazilian College of Animal Experimentation (COBEA) and approved by the Institutional Ethics Committee (CEUA/UNICAMP #3655-1) were followed throughout the study. Our local colonies originated from breeding stock supplied by CEMIB/UNICAMP, Campinas, SP, Brazil. Immediately after weaning at 3 weeks of age, animals were maintained under controlled temperature (25°C) and lighting conditions (08:00–18:00 h), with *ad libitum* access to tap water and standard rodent laboratory chow (Nuvital, Curitiba, PR, Brazil); rats were raised up to 12 weeks of age.

Dams were maintained in an isocaloric standard rodent laboratory with *ad libitum* access to either regular protein content (NP) (17% protein, n = 20) or low protein content (LP) (6% protein, n = 20) chow for the duration of their pregnancy (Table 1).

**Table 1.**
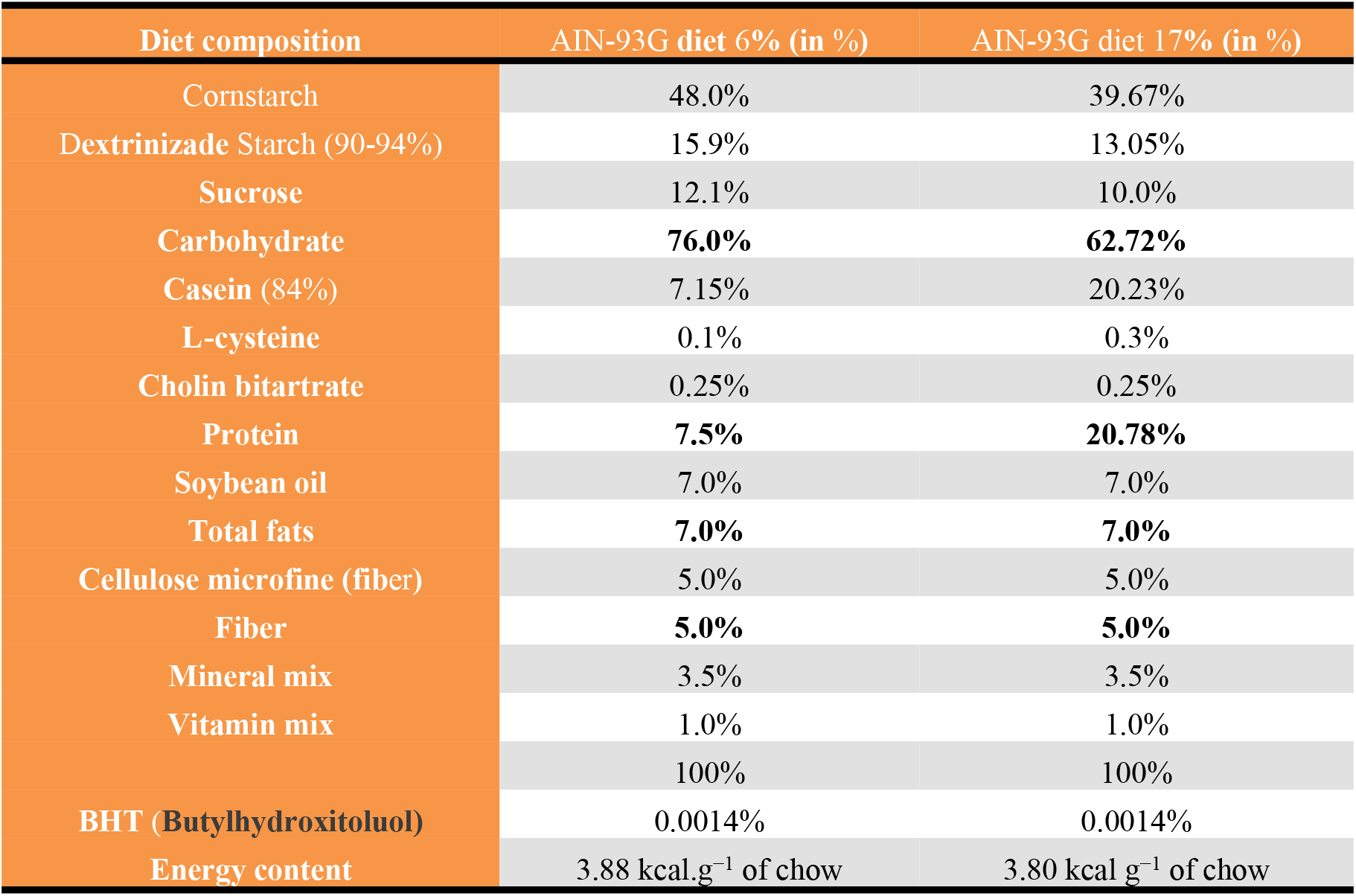
Composition of standard rodent laboratory diet (standard normal-protein (NP) diet, 17% and low-protein (LP) diet, 6% (AIN 93G)

At the time rats were placed to mate, sperm in the vaginal smear was designated as day 1 of pregnancy. Dams were kept in individual metabolic cages and weighed during pregnancy. Bodyweight, food consumption, and protein consumption were determined daily, and from these values, the weekly average was recorded once a week for each pregnant dam. Protein intake for each week was calculated by considering the total food intake and protein content of each diet.

On the day of birth, male offspring were weighed; only eight pups were kept per female. All groups returned to NP chow intake after delivery. We studied male LP offspring between 48 and 88 weeks of age, comparing them to age-matched NP offspring for analysis. An animals’ survival lifetime in both groups (NP; n = 21 and LP; n = 18) was established as the spontaneous death of offspring in their previously defined housing.

### 2. Behavioral analysis

Male offspring from each litter, NP (n=20) and LP (n=20) groups, were used for behavioral research. All behavioral tests were performed during the light cycle. Test room illumination was kept constant and controlled under low-intensity white lighting (5–30 lux).

#### 2.a. Morris water maze (MWM)

All-cognitive performance analysis was conducted using the MWM test, a spatial learning test for rodents [22]. The test was performed with NP and LP offspring at 48 and 50 weeks and between 86 and 88 weeks of age (n = 20 for each group) by a trained observer in a blinded fashion, as previously described [22]. Behavioral tests were performed in a circular black tank (170-cm diameter) filled to a depth of 31 cm at 22°C and placed in a dimly lit room with extrinsic clues. The tank was divided into imaginary quadrants, with a black platform (12-cm diameter, 30-cm height) placed in one.

#### 2.b. Working memory task

A test described by Kesner et al. (2000) was used to test prefrontal cortex (PFC) function. This test aimed to assess the ability of a rat to learn the position of a hidden platform and to retain this information during four consecutive trials [23]. The working memory test consisted of 4 days of acquisition, with four trials per day. On each trial day, the position of the platform was kept constant; however, its position was varied on each successive day such that all four quadrants were used. Rats were placed facing the wall of the maze at a different starting point, north (N), east (E), south (S), or west (W), at the start of each daily trial. A trial was considered complete when the rat escaped onto the platform. When this escape failed to occur within 120 s, the animal was gently guided to the platform. Rats were allowed to spend 30 s on the escape platform before being positioned at a new starting point. Escape latency, defined as the time (in seconds) spent to reach the platform, was recorded in each consecutive trial and expressed as the mean of each trial day.

#### 2.c. Reference memory task

Morris (1984) described the reference memory test used to assess hippocampal function. This test aimed to evaluate the ability of a rat to learn the position of a hidden platform and retain this information during all test days [22]. The test consisted of 4 days of acquisition, with four trials per day. During the 4 days, the platform remained in the same quadrant. Rats were placed facing the wall of the maze at a different starting point, N, E, S, or W, at the start of each of the four daily trials. The trial ended when the rat escaped onto the platform. When this escape failed to occur within 120 s, the animal was gently guided to the platform. Rats were allowed to spend 30 s on the escape platform before being positioned at a new starting point. The representative steeper curve of the time (in seconds) spent to reach the platform (escape latency) was recorded in consecutive trials and expressed as the mean of each trial day.

#### 2.d. MWM proximity measure

The test was performed in LP (n=10) and age-matched NP (n=10) offspring between 86 and 88 weeks of age. The animals were trained using a standardized procedure that required distal cues in the maze environment to learn the position of a hidden escape platform [24]. Briefly, rats were tested for four trials per day, with inter-trial delays of 30 sec, for a total of four consecutive days. The platform’s location remained constant, and on each training trial, animals swam for 120 sec or until they found the platform. Across trials, the starting location varied among four equidistant points around the perimeter of the apparatus. Every fourth trial was a probe test during which the platform was retracted to the pool’s bottom. Probe trial performance provides an assessment of the search strategies used to navigate the maze [24]. Spatial behavior was recorded throughout training using a video tracking system designed for this purpose (Institute of Computing, Unicamp, SP, Brazil). The status of spatial learning was evaluated during each probe trial. The rat’s distance from the escape platform was sampled 10 times per sec, and these values were averaged in 1-sec bins. Cumulative search error was then calculated as the summed 1-sec averages of this proximity measure corrected for each trial’s particular start location (in mm). This measure reflects the average proximity during probe trials relative to the escape platform’s training location; lower index scores indicate a more accurate search focused on the target location. Previous research demonstrates that these measures are especially useful for distinguishing spatial learning impairments from other age-related performance deficits that are presumably independent of hippocampal function [24].

#### 2.e. Elevated plus maze (EPM)

Next, 48 to 50 and 86 to 88-week-old LP and NP offspring (n = 20) were tested for 5 min in an EPM, a 72-cm high white polypropylene “plus”-shaped maze (ENV-560; Insight Equipment Ltd, Ribeirão Preto, SP, Brazil). The maze consisted of two facing open arms (50.8 × 10.2 cm) and two closed arms (50.8 × 10.2 × 40.6 cm). The time spent in the open arms, junction area, and closed arms and the number of entrances and explorations in each section, were recorded using a system of infrared photo beams, with the crossings monitored using a computer. The duration spent in each EPM compartment was presented as a percentage of the trial’s total duration.

### 3. Isotropic fractionator study

Hippocampal cells from NP and LP offspring were quantified using a technique previously described by Herculano-Houzel and Lent (2005) [25]. Briefly, following behavioral tests, sacrificed 88 week-old NP (n = 5) and age-matched LP (n = 5) offspring were perfused with saline, followed by 4% buffered paraformaldehyde, using cardiac puncture. Next, hippocampi were rapidly dissected on ice under a stereomicroscope; anatomical landmarks were observed according to Paxinos and Watson (2005) using a standardized technique widely used by the Life and Health Sciences Research Institute (ICVS), School of Health Sciences, University of Minho, Braga, Portugal. Following isolation of the left hippocampus, a suspension of nuclei was obtained through mechanical dissociation in a standard solution (40 mM sodium citrate and 1% Triton X-100) in a 40-mL glass Tenbroeck tissue homogenizer.

Complete homogenization was achieved using at least 1 mL of dissociation solution per 100 mg of brain tissue and by grinding until the smallest visible fragments were dissolved. The homogenate was collected using a Pasteur pipette and transferred to 15-mL centrifuge tubes. The grinding pestle and tube were washed several times with dissociation solution, and the sample was centrifuged for 10 min at 4000× g. Pelleted nuclei were then suspended in phosphate-buffered saline (PBS) containing 1% 4’,6-diamidino-2-phenylindole (DAPI) (Molecular Probes, Eugene, OR, USA) to make all nuclei visible under ultraviolet illumination. After sufficient agitation, 5-μL aliquots were used to determine nuclei density using a hemocytometer. The DAPI-stained nuclei were counted under a fluorescence Olympus microscope at 400× magnification.

Once the nuclear density in the suspension was determined by averaging at least eight samples, the total number of cells in the original tissue was estimated by multiplying the mean nuclear density by the total suspension volume.

To estimate the total neuron number, a 200–500-μL aliquot of nuclear suspension was immunoreacted for NeuN. Nuclei in the aliquot were collected using centrifugation, resuspended in a 0.2 M solution of boric acid (pH 9.0), and heated for 1 h at 75°C for epitope retrieval. Subsequently, nuclei were again collected using 6000xg centrifugation, washed in PBS, and incubated overnight at room temperature with anti-NeuN mouse antibody (1:300 in PBS; Chemicon, Temecula, CA, USA).

The nuclei pool was washed in PBS and incubated in cyanine 3-conjugated goat anti-mouse IgG secondary antibody (1:400 in 40% PBS, 10% goat serum, and 50% DAPI; Accurate Chemicals, Westbury, NY, USA) for 2 h. Next, the centrifugation-collected nuclei were washed and resuspended in PBS prior to counting under a fluorescence microscope. The total number of non-neuronal nuclei was calculated by subtracting the number of NeuN-containing nuclei from the total number of nuclei.

### 4. Immunoblotting

Eighty-eight-week-old NP (n = 5) and age-matched LP (n = 5) offspring were anesthetized and their hippocampus isolated in ice-cold buffer for analysis. The Bradford method was used for protein quantification in tissue extracts. For immunoblotting of total protein, extracts were loaded onto an SDS-PAGE electrophoresis gel. Briefly, equal amounts of protein (20 μg) from hippocampal samples were denatured in Laemmli buffer (100 mmol/L DTT), incubated at 95°C for 4 min, and electrophoresed on an 8% SDS-PAGE gel. Proteins were transferred from the gel to nitrocellulose membranes for 90 min at 120 V (constant) using a Bio-Rad miniature transfer apparatus (Mini-Protean, Bio-Rad).

Non-specific protein binding to the nitrocellulose was reduced by preincubating the filter for 2 h at 22°C in blocking buffer (5% non-fat dry milk, 10 mmol/L Tris, 150 mmol/L NaCl, and 0.02% Tween 20) [26]. Nitrocellulose blots were incubated at 4°C overnight with specific antibodies against IGFR1 (Ab90657; Abcam, Cambridge, UK, dilution, 1:1000), ERK2 (sc154; Santa Cruz Biotech, Inc., Santa Cruz, CA, USA, dilution, 1:1000), pPI3K (cs4228S; Cell Signaling Technology, Danvers, MA, USA, dilution, 1:1000), brain-derived neurotrophic factor (BDNF) (sc546; Santa Cruz Biotech, Inc., dilution, 1:1000), Tau (sc5587; Santa Cruz Biotech, Inc., dilution, 1:1000), pTau (sc16945; Santa Cruz Biotech, Inc., dilution, 1:1000), Aβ (sc 374527; Santa Cruz Biotech, Inc., dilution, 1:1000), HSP70 (sc3575; Santa Cruz Biotech, Inc., dilution, 1:1000), and HSP90 (sc7947; Santa Cruz Biotech, Inc., dilution, 1:1000).

Membranes were subsequently incubated with peroxidase-conjugated secondary antibodies (1:10.000). Bands were detected using chemiluminescence (RPN 2108 ECL; Amersham Pharmacia Biotech, Piscataway, NJ, USA); density was quantified using the optical method (Scion Image 4.0.3.2 software; ScionCorp, Frederick, MD, USA). Images of developed radiographs were scanned (Epson Stylus 3500), and band intensities quantified using optical densitometry (Scion Image Corporation). Membranes were stained with α-tubulin (cs2144S; Cell Signaling Technology, dilution, 1:1000) or reversible Ponceau to discard any possible inequalities in protein loading and transfer in the western blots. Only homogeneously stained membranes were used in this study.

### 5. Data presentation and statistical analysis

All data were reported as mean ± standard error of the mean (SEM). Data obtained over time were analyzed using one-way analysis of variance (ANOVA). When one-way ANOVA analysis indicated statistical differences between groups, post-hoc comparisons between means were performed using Bonferroni’s contrast test. Comparisons between two groups were performed using a two-way repeated measure ANOVA, in which the first factor was protein content in a dam’s diet, and the second factor was time. When the interaction was significant, mean values were compared using Tukey’s post-hoc analysis. Student’s *t*-tests were used to evaluate studies involving only two independent samples, within or between groups. Welch’s test was used to correct situations characterized by heteroscedasticity (different variances between groups). An animal’s survival lifetime was assessed using Mantel-Cox and Gehan-Breslow-Wilcoxon tests. GraphPad Prism 5.00 was used for data analysis (GraphPad Software, Inc., La Jolla, CA, USA). The level of significance was set at p ≤ 0.05.

## RESULTS

### 1. A gestational low-protein diet affects birthweight and the hippocampal mass

Although body mass of LP male pups at birth was significantly reduced, compared to NP progeny (NP: 6 ± 0.05 g, n = 20 vs. LP: 5.8 ± 0.05 g, n = 20, p = 0.006, Figures 1A and S1); after 7 days of delivery, the LP and NP offspring body mass are similar, and this similarity persisted unchanged until week 88 of follow-up. (Figure 1C-E). Brain and hippocampal masses did not differ between 50-week-old LP (n = 5) and NP offspring (n = 5, Figure S2). However, in 88-week-old LP, the relative hippocampus body (H/bd) or brain (H/br) mass ratios were significantly enhanced relative to an age-matched NP hippocampal mass (H/bd, LP: 0.03 ± 0.004 vs. NP: 0.01 ± 0.001, n = 5, p = 0.03 and H/br, LP: 0.06 ± 0.005 vs. NP: 0.04 ± 0.006, n = 5, p = 0.01, Figure 1C-E, bottom Panels). Dams on the LP diet during pregnancy were lighter in the second (238.13 ± 6.0 g) and third (325.1 ± 6.6 g) weeks of pregnancy as compared to those on the NP (2nd week: 244.7 ± 8 g and 3rd week: 354.7 ± 8 g) diet, despite having an equal body weight in the first week of pregnancy (diet vs. time p < 0.001; diet p = 0.007; time p < 0.001). Thus, considering the entire pregnancy, dams in the LP group had lower weight gain than those in the NP group (NP [n = 20]: 109.86 ± 20.04 g; LP [n = 20]: 87.00 ± 14.01 g; p < 0.001). Weekly food intake was higher in LP dams in the first two weeks of pregnancy (1st week: 129.2 ± 3.25 g and 2nd week: 135.7 ± 6 g) compared to NP (1st week: 111.1 ± 3.95 g and 2nd week: 119.4 ± 6 g) dams (diet vs. time p = 0.018; diet p < 0.001; time p < 0.118). However, there was no difference in food intake between the groups in the last week of pregnancy (NP: 122.94 ± 3.95 g vs. LP: 122.9 ± 6 g). Litter sizes were similar (p = 0.239) between rats fed an LP diet (n = 10.67 ± 2.44 puppies) and those fed an NP diet (n = 11.43 ± 1.94 puppies). The weekly assessment of protein intake showed that dams from the LP group were exposed to severe protein intake restriction (NP: 20.01 ± 0.9 g vs. LP: 7.91 ± 0.7 g) during the entire pregnancy (diet vs. time p = 0.018; diet p < 0.001; time p = 0.069).

**Figure 1.**
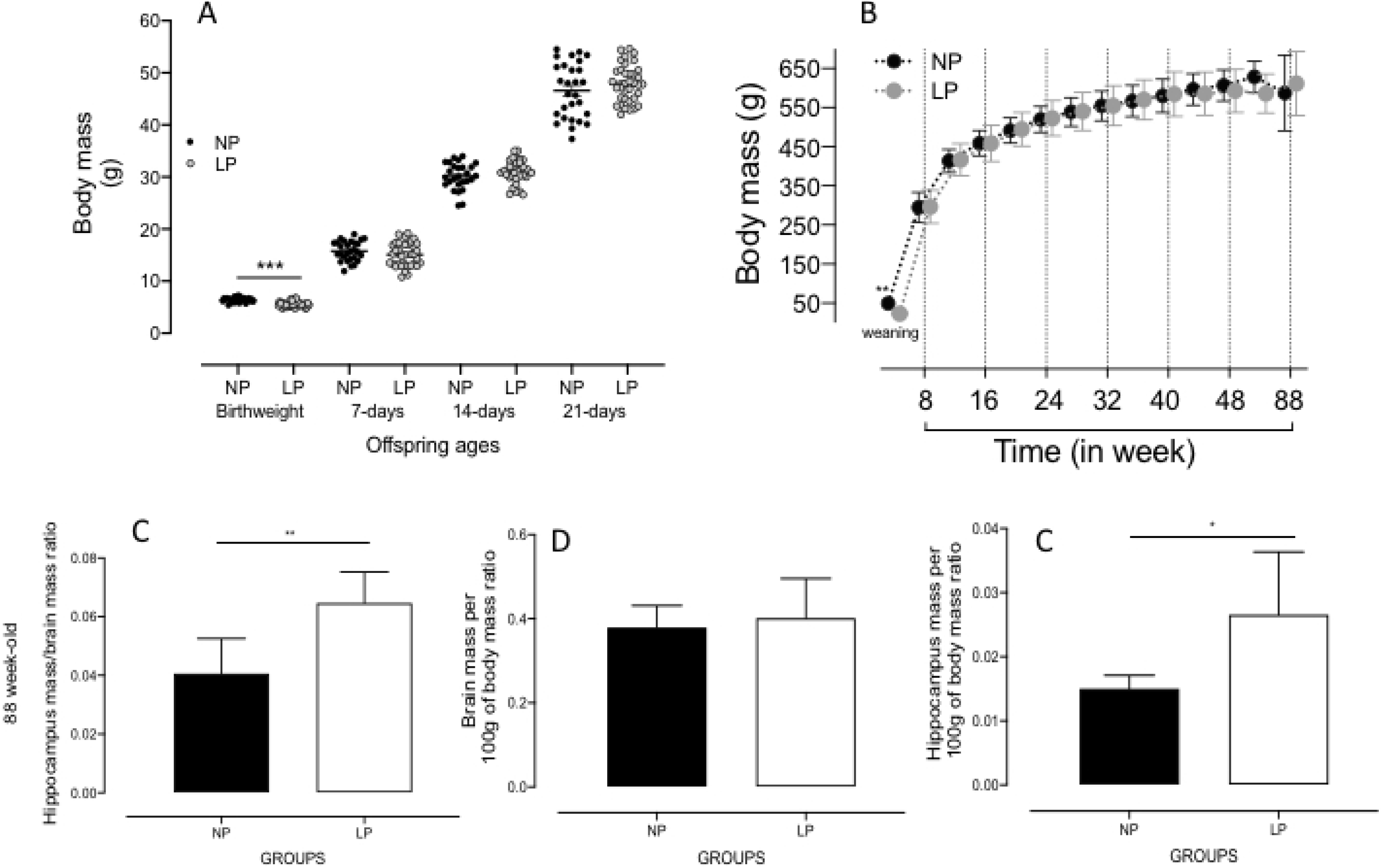
(A) Offspring bodyweight (g) at birth to 21 days of age, and (B) in 4 to 88-week-old NP (n=20), as compared to LP (n=20) offspring. Also, the figure shows the offspring brain and hippocampal masses (g) in 88-week-old NP (n = 5) as compared to age-matched LP (n = 5) offspring (Figure 1 C-E, Bottom Panels). Results are expressed as means ± SEM; data obtained over time were analyzed using repeated-measures ANOVA, and comparisons involving only two means within or between groups were performed using a Student’s *t*-test. Welch’s *t*-test was used to correct situations characterized by heteroscedasticity. When statistically significant differences were indicated between selected means, posthoc comparisons were performed using Bonferroni’s contrast test. The level of significance was set at *p < 0.05.

### 2. Morris Water Maze (MWM)

#### 2.a. Working memory

Working memory, as estimated via escape latency, was not statistically different in 46 and 86-week-old LP as compared to age-matched NP offspring (49-week old, n = 20, p = 0.87, or 87-week old, n = 20, p = 0.078) (Figure 2). These results suggest the same learning response in both experimental groups.

**Figure 2.**
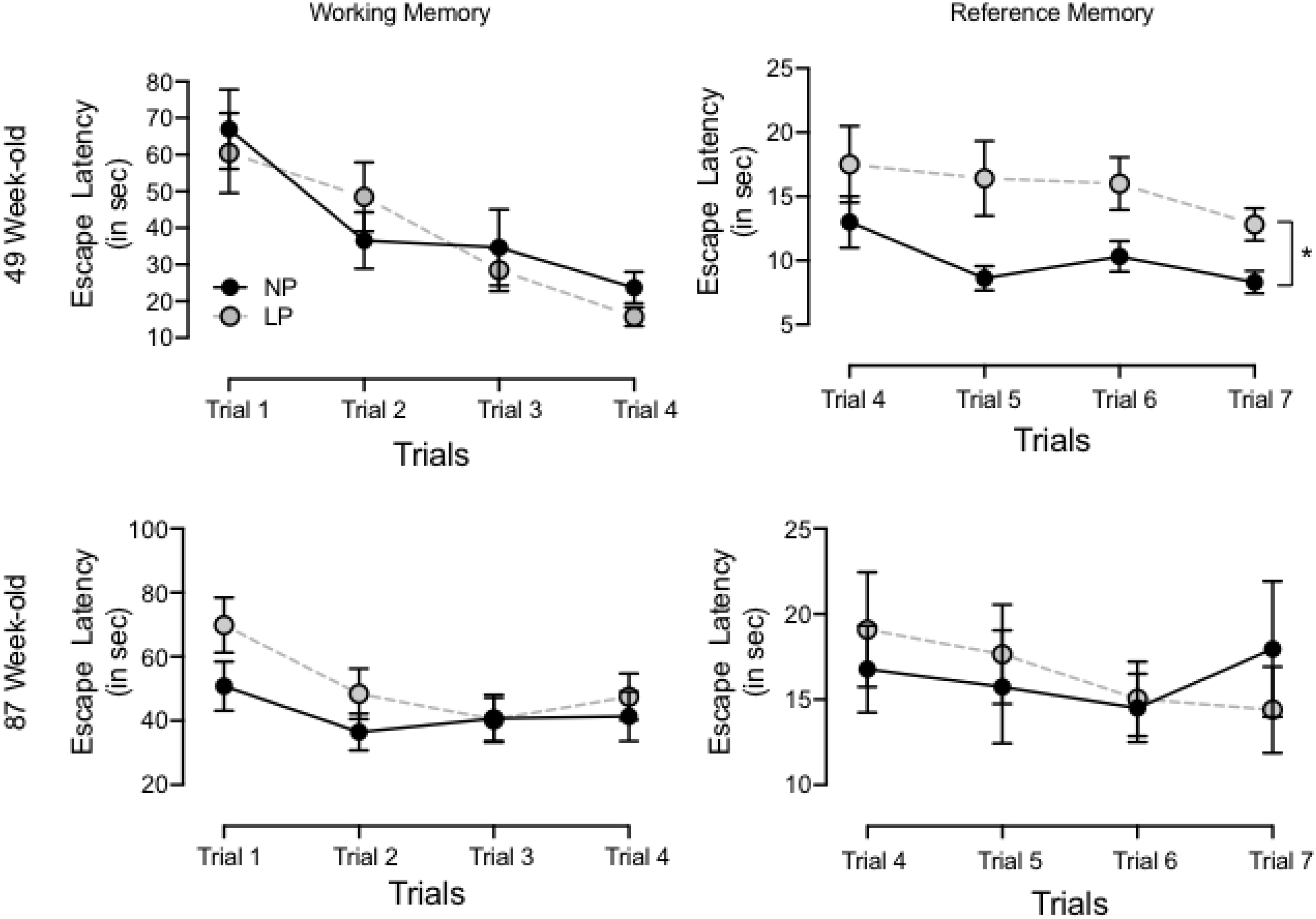
Graphic representation of the Morris water maze test was performed with 46 and 86-week-old LP (n = 20) compared to age-matched NP (n = 20) offspring to assess both working and reference memory. The working memory test consisted of 4 days of acquisition (4 trials/day). The time (seconds) spent to reach the platform (escape latency) was recorded in consecutive trials and expressed as each trial day’s mean. Results are expressed as means ± SEM; data obtained over time were analyzed using repeated-measures ANOVA. When statistically significant differences were indicated between selected means, posthoc comparisons were performed using Bonferroni’s contrast test. The level of significance was set at *p < 0.05.

#### 2.b. Reference memory

Hippocampal reference memory significantly reduced in 49-week-old LP rats as compared to NP offspring (p = 0.0372). These animals spent substantially more time finding the hidden platform (latency) than their age-matched NP offspring. When comparing day 4 and 7 trials, reduced latency (learning capacity) was observed only in NP (n = 20, p = 0.046) as compared to LP offspring (n = 20, p = 0.08, Figure 2). However, in the 87-week-old offspring analysis, no reference memory differences were observed in either group.

#### 2.c. Proximity measure

The rat’s distance from the escape platform was sampled 10 times per sec, and these values were averaged in 1-sec bins. An elevated cumulative search error, calculated as the summed 1-sec averages of this proximity measure corrected for each trial’s particular start location (in mm), was observed in 87-week-old LP as compared to NP rats (LP: 52.6 × 10^4^ ± 10.3 × 10^4^ mm *vs.* NP: 47.0 × 10^4^ ± 10.6 × 10^3^ mm, p = 0.0007; n = 10 for each group, Figure. 3).

**Figure 3.**
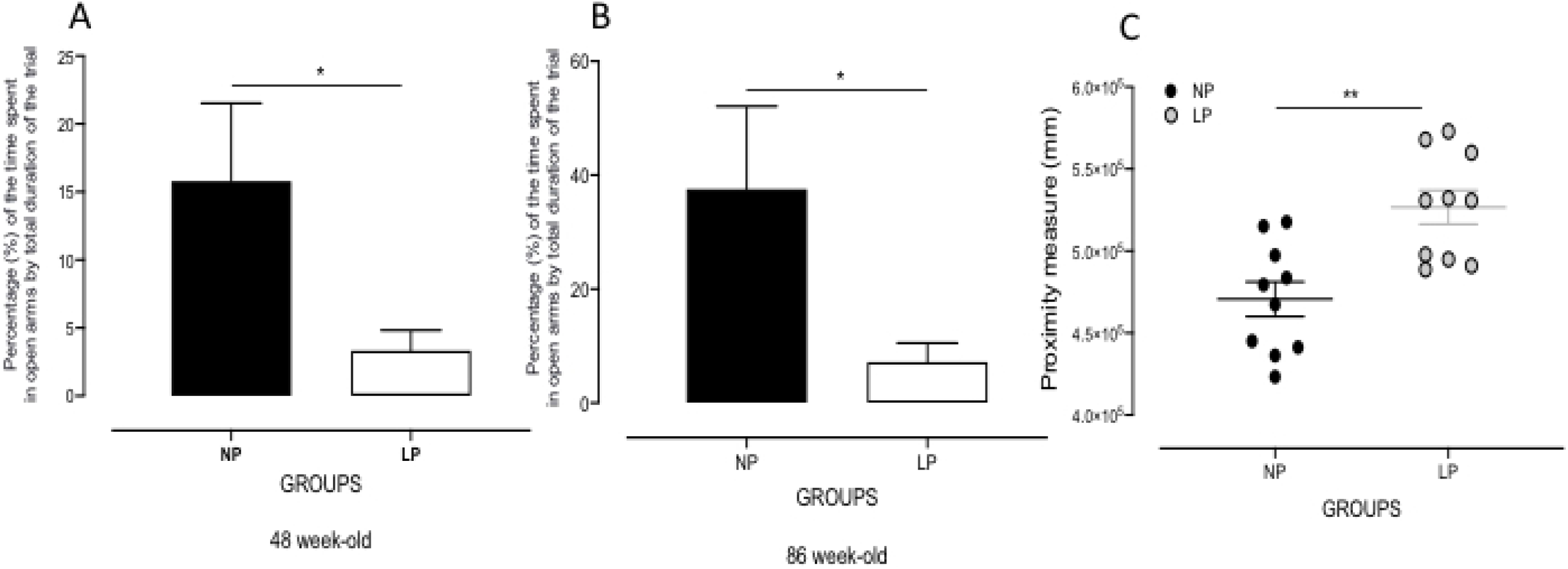
Graphic representation of the elevated plus maze (EPM) test performed with 46 and 86-week-old LP (n = 9) as compared to age-matched NP (n = 8) offspring (A and B Panels). Values of time spent in EPM open arm compartments are presented as percentages of the trial’s total duration. The C Panel shows the proximity measure test’s graphic representation performed with 86-week-old LP (n = 8) compared to age-matched NP (n = 8) offspring. Results are expressed as means ± SEM; comparisons involving only two means within or between groups were performed using a Student’s *t*-test. The level of significance was set at *p < 0.05.

### 3. EPM

The gestational protein-restricted diet triggered anxiety-related behaviors in adult offspring as compared to NP adult progeny at both 48 and 86 weeks of life. The time spent in each EPM compartment was presented as a percentage of the trial’s total duration. The results showed a significant reduction in the time spent in open arms (in %) in 48 (NP: 15.78 ± 5.749% *vs*. LP: 3.321 ± 1.512%, p = 0.0348, n = 8) and 86 (NP: 37.55 ± 14.55% *vs*. LP: 7.171 ± 3.357%, p = 0.0388, n = 8, Figure 3) week-old LP compared with age-matched NP offspring.

### 4. Quantification of total cells and neurons in the hippocampus

Total cell number was significantly enhanced in the hippocampus of LP as compared to NP rats (LP: 15.98 × 10^6^ ± 17.69 × 10^5^ *vs.* NP: 12.56 × 10^6^ ± 33.85 × 10^4^, p = 0.047, n = 8 for each group, Figure 6). The estimated number of neurons was unaltered in LP as compared to NP offspring. However, glial cell numbers were enhanced in LP as compared to NP offspring (LP: 13.91 × 10^6^ ± 11.66 × 10^6^ *vs*. NP: 10.58 × 10^6^ ± 30.03 × 10^4^, p = 0.04, Figure 4).

**Figure 4.**
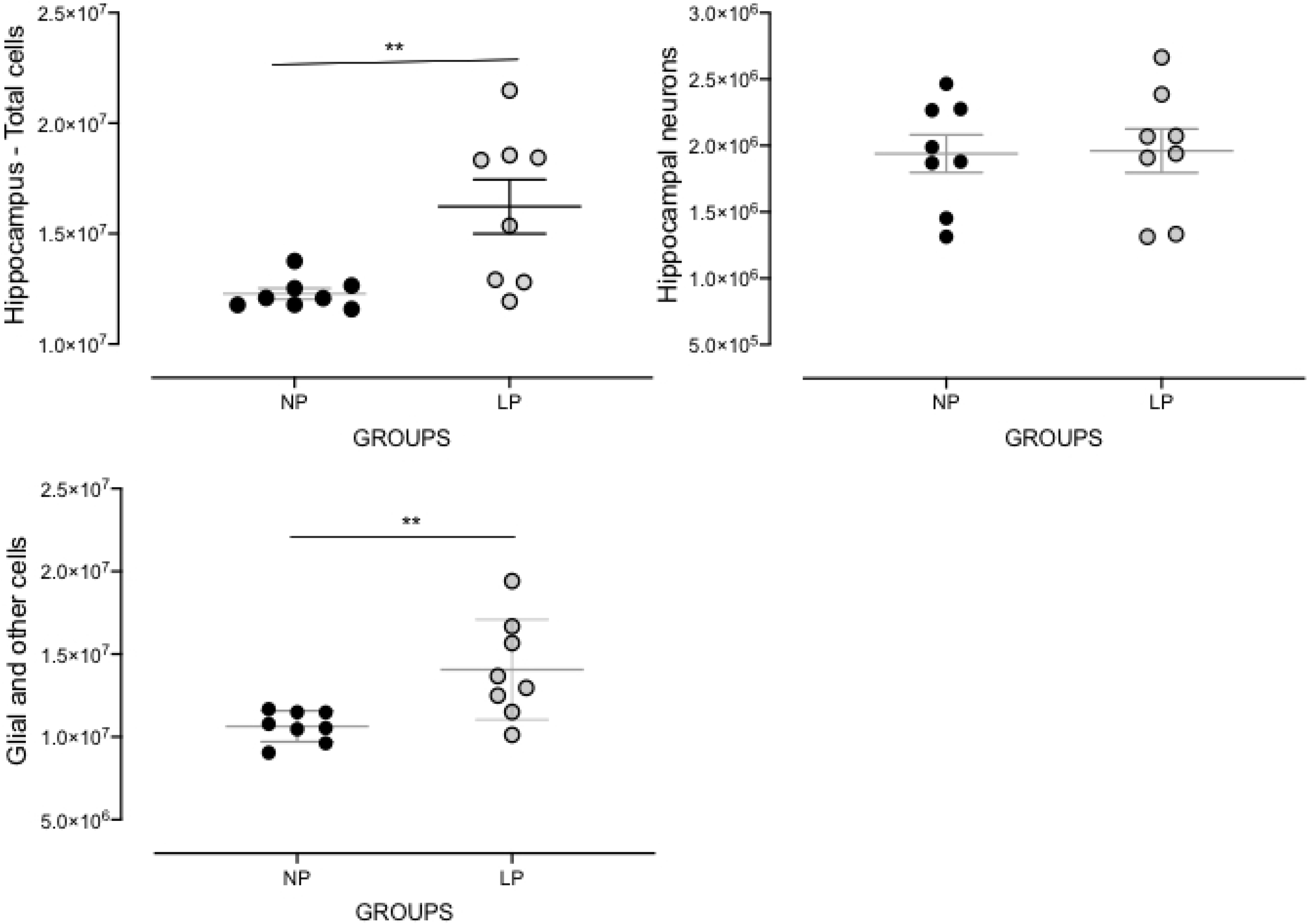
Effects of maternal protein restriction on 88-week-old LP (n = 5) compared to age-matched NP (n = 5) offspring on whole hippocampal cell, neurons, and non-neuronal cell quantifications. Results are depicted as scatter dot-plot and are expressed as means ± SEM; comparisons involving only two means within or between groups were performed using a Student’s *t*-test. The level of significance was set at *p < 0.05.

### 5. Immunoblotting

Whole hippocampal protein levels were assessed in 88 week-old NP compared to age-matched LP offspring using immunoblotting comparative analysis. The results revealed 61% enhanced BDNF mature protein levels in LP compared to NP offspring (n = 5 for each group, p = 0.0413; Figure 5). Similarly, whole hippocampal extracts showed increased tau, tau phosphorylation (48%, n = 5, p = 0.0397), and Aβ (77%, n = 5, p = 0.0365) protein levels in LP compared to age-matched NP offspring. However, stress-related proteins (HSP70 and HSP90), IGFR1, pPI3K, and ERK2 were unchanged between both groups (Figure 5).

**Figure 5.**
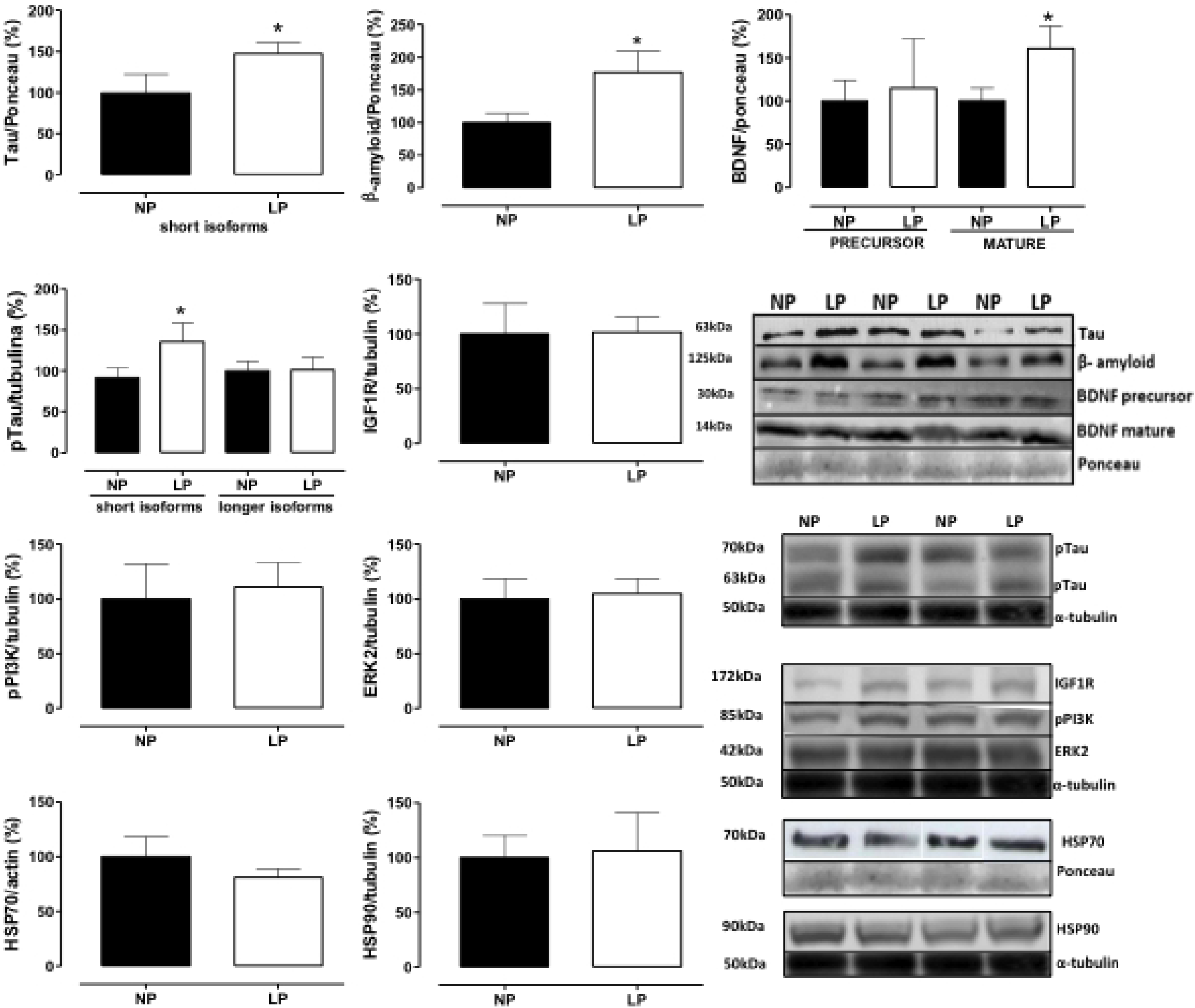
Effect of maternal protein restriction on β-amyloid, tau, BDNF, HSP70, HSP90, IGF1R, pPI3K, and ERK2 proteins, measured using immunoblotting analysis, in the isolated whole hippocampus of 88-week-old LP (n = 4) as compared to age-matched NP (n = 4) rats. Results are expressed as means ± SEM; only one offspring of each litter was used for immunoblotting experiments. A comparison involving only two samples of independent observations was performed using a Student’s *t*-test. The level of significance was set at *p < 0.05.

### 6. Survival curve

The survival lifetime of animals, as evaluated using Mantel-Cox and Gehan-Breslow-Wilcoxon tests, demonstrated a significant reduction (p < 0.001) in the life span of LP (n = 18) as compared to NP rats (n = 21). The estimated median survival time of NP and LP rats was 120 weeks and 108 weeks, respectively. These results showed a significant reduction in survival time in LP as compared to NP offspring (Figure 6).

**Figure 6.**
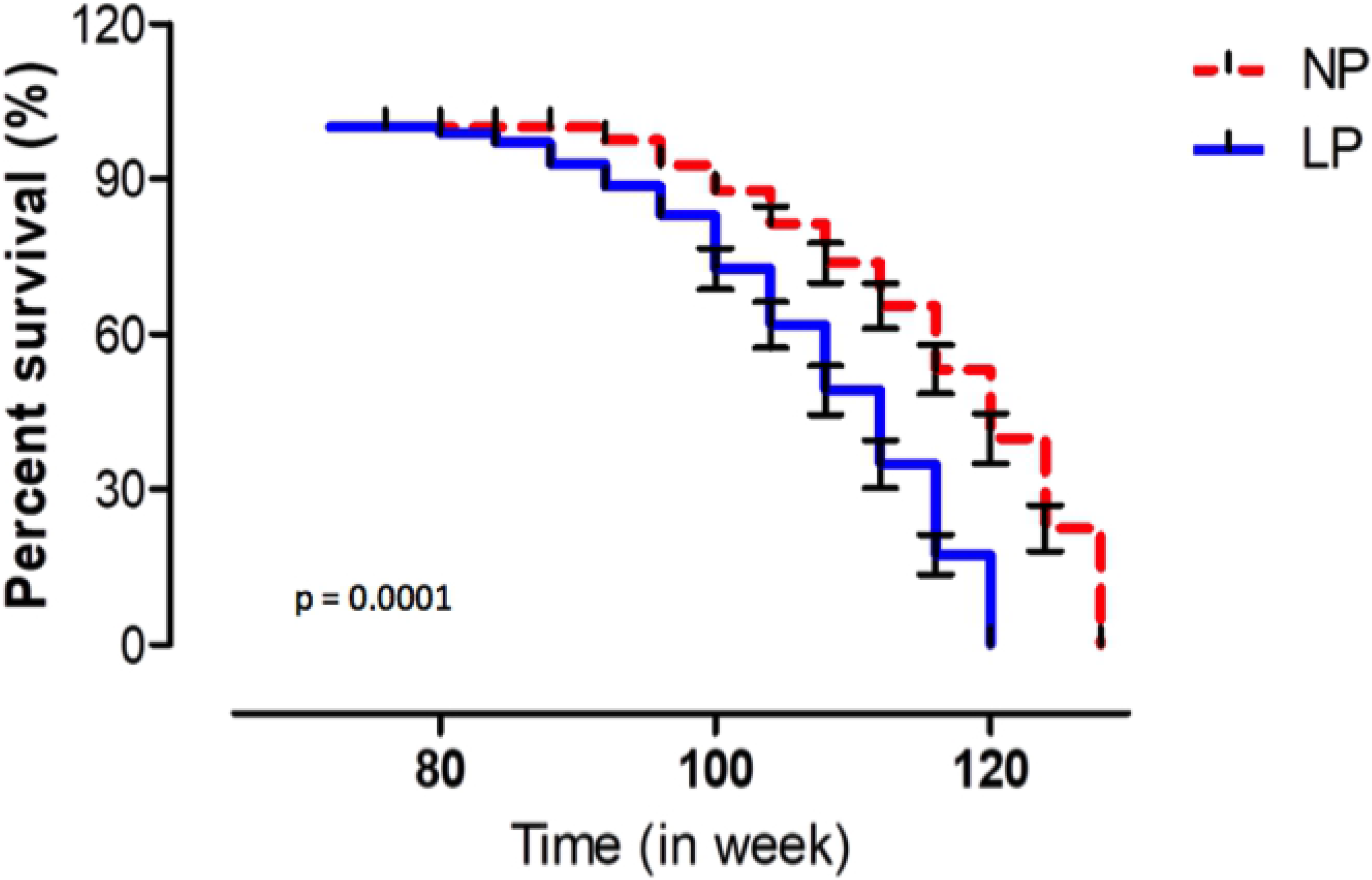
Survival curve evaluated using Mantel-Cox and Gehan-Breslow-Wilcoxon tests in NP (n = 10) compared to LP (n = 10) offspring. Results are expressed as means ± SEM; the significance level was set at *p < 0.05.

## DISCUSSION

Studies have demonstrated that disturbances during crucial fetal developmental periods may determine morphological and functional changes in organs and systems [1–6]. Brain development is sensitive to gestational programming, and humoral factors, such as angiotensin, steroids and their receptors, transcriptional factors, and nutritional availability, permanently affect neural ontogenesis [27,28]. Among neural structures, the hippocampus has been demonstrated to be particularly susceptible to the programming effects of gestational emotional and nutritional stressors [for a review, see 29].

The present study confirms previous data demonstrating that gestational protein restriction leads to low birth weight and catch-up growth phenomena in rodents. This effect leads to gender-related disorders in blood pressure, glucose metabolism, and anxiety-like behaviors in male adult progeny compared to female offspring [30,31]. Sex hormones determine phenotypic dimorphism in an adult fetal-programmed disease model by modulating regulatory pathways critical for the long-term control of neural, cardiac, and endocrine functions.

Specific hormones and the estrus cycle may interfere with behavioral, hemodynamic, and systemic water and ion homeostasis in female rodents. Thus, considering the above findings and the influence of gender on physiological responses, this study was performed in male rats. However, additional research with long-term follow-up and cross-fostering, including behavioral tests in female offspring, would help specify the nature of some protein-restriction effects. Our previous results sought to elucidate the underlying mechanisms of fetal programming [11,12,15,32–37]. The present study confirms that reduced fetal birth weight in LP offspring may reflect the influence of inappropriate protein intake on healthy embryo growth and development during pregnancy.

As mentioned above, after the second week of age, LP offspring’s body mass was similar to that observed in the NP group, a phenomenon known as catch-up growth. Also, the brain weights of 50 and 88-week-old LP rats were the same as that for age-matched NP offspring. However, in 88-week-old LP rats, the hippocampus weight by body or brain mass ratio was significantly enhanced as compared to age-matched NP hippocampal ratio masses. Additionally, an 88-week-old LP hippocampi cell study using the isotropic fractionator technique showed an increased number of glial and other non-neuronal cells; the neuron number was not otherwise altered as compared to age-matched NP offspring.

Previous studies investigating the effects of maternal undernutrition on brain structure have demonstrated loss of neurons [14], decreased neuronal body size and length of apical dendrites, branching [15,35,38], and reduced CA1 hippocampal subfield pyramidal cell spine density [39]. These studies have also shown dentate gyrus and decreases in the somatic axis and the number of dendritic branches and spines [15,38,40]. However, to the best of our knowledge, no predicted studies have evaluated changes in the hippocampal number and type of cells in severe maternal protein-restricted offspring.

Previous studies have demonstrated that several functional neural disorders may result from substantial glial proliferation, which can occur to differing extents in specific brain regions [41]. As reactive gliosis is a hallmark of AD, we may infer that an increased number of hippocampal glial cells may be associated with structural and functional disorders in this neurodegenerative rodent model [42,43]. Herein, hippocampi homogenate was processed following vigorous whole-brain saline perfusion to minimize contamination with peripheral cells. However, we cannot exclude a potential contribution from infiltrating cells once the blood-brain barrier is compromised during Alzheimer-like disease [44–46].

We hypothesize that hippocampi with enhanced glial cells may have potential roles in the spread of tau pathology [47], synapse loss [48], and neuronal dysfunction through the release of pro-inflammatory factors [49,50]. In 1907, Alzheimer himself described reactive glial cells in neuritic plaques [51]. A recent study confirmed the presence of reactive astrocytes and microglia in the vicinity of Aβ deposits [52], both of which are associated with neuroinflammation and AD evolution. Although the role of glial cell participation is unclear, the high number of hippocampal non-neuronal cells strongly suggests a causal nexus despite conflicting reports regarding their detrimental or protective contribution to AD pathogenesis [53].

However, emerging studies report a loss of oligodendrocyte lineage cells in gray matter, presumably associated with amyloid plaque deposition, oxidative stress, and apoptosis [54–56]. An accurate understanding of the role of astrocytes in AD remains unclear. Considering the present model’s findings, we hypothesize that glial cells may play a critical role in dementia processes, likely through microglial activation. Further studies on cellular composition are essential to better understand hippocampal contributions to pathological behavior manifestations.

The present study evaluated whether gestational protein-restricted intake is associated with hippocampal molecular changes and, if so, aimed to explain the functional disorder in aging male LP progeny. In addition, to the best of our knowledge, this study is the first to show a significant enhancement of hippocampal Aβ deposition in maternal protein-restricted 88-wk-old offspring.

Although increased tau and Aβ deposition was consistently demonstrated in the 88-week-old LP hippocampus as compared to age-matched NP offspring, stress-related proteins, HSP70 and HSP90, were unchanged in both groups. Typical AD-associated structural brain abnormalities are characterized by senile plaques and NFT, which are complexes of paired helical microtubule-protein tau filaments associated with toxic Aβ oligomer deposition, resulting from Aβ monomer self-association [57–59]. Prior research has demonstrated that Aβ monomers are naturally produced and secreted at the synaptic cleft; Aβ monomers appear to be necessary for neuronal survival, given that Aβ production inhibition is associated with cultured neuronal death [60].

The present study showed that, despite enhanced hippocampal deposition of Aβ oligomers in maternal protein-restricted adult offspring, the hippocampus presented a significant increase in BDNF expression. BDNF is significant for maintaining the development and survival of neurons in the brain. Considering primary data, BDNF may be essential for maintaining viable cortical neurons, whose early dysfunction contributes to short-term memory loss [61].

However, if and how glial dysfunction is involved in changes to hippocampal circuitry and its contribution to learning and memory disorders remain unknown. Herein, using the MWM test, we assessed learning and memory in gestational protein-restricted and NP progeny, finding significant differences in working memory responses, as characterized by enhanced escape latency in 49-week-old LP as compared to age-matched NP offspring, but not in older offspring.

However, a highly sensitive assessment of age-related learning/memory impairment in aged rats was performed using proximity measure distances [24], which increased in 86-week-old LP, as compared to age-matched NP offspring. In the current study, proximity measures were useful for associative analyses, taking into account changes in hippocampal neurocytology, abnormal protein deposition, and LP offspring’s functional decline. These data suggest that the hippocampal referential memory process in LP offspring might be altered by maternal protein restriction, particularly in elderly rats.

Also, LP offspring were found to spend less time in EPM open arm compartments, resulting in an extended stay in the device’s closed arms, as presented via percentages of the trial’s total duration.

These results revealed an anxiety-like behavioral phenotype in 88-week-old LP as compared to age-matched NP offspring. The current study sought to explore the relationship between elevated hippocampal mass, potentially via increases in glial and other non-neuronal cells, and anxiety-like behavior in 88-week-old LP rat progeny. Conversely, in human studies, patients with depression and previous depression-induced animal models have demonstrated a negative relationship between hippocampal volume/mass and trait anxiety [62,63].

However, the present study was unable to exclude the contribution of several brain regions in rodent anxiety-like behavior as well as possible confounds arising from the segregation of other behavioral disorders. Considering the above findings, we suppose that enhanced BDNF in 88-week-old LP offspring is a final attempt to maintain hippocampal functionality and protect it from further damage. Within this scenario, elevated hippocampal BDNF levels may restrain neuron loss, at least temporarily, even reducing partially hippocampal dysfunction previously altered in 49-week-old LP offspring.

Preliminary data have shown the involvement of hippocampal molecular pathways, such as PI3K/AKT, by recruiting IGF-IRs and increasing CREB expression via Aβ monomer stimulation during increased transcription and release of BDNF [63–65]. We could not demonstrate any change in IGF-IR and PI3K phosphorylation in the current study, suggesting that BDNF may be expressed via different mechanisms [53]. Recent studies have also indicated that tau protein is a crucial factor underlying AD development and progression. The present model confirmed such findings, with increased hippocampal tau expression accompanying 28% enhancement in phosphorylated tau in 88-week-old LP as compared to age-matched NP rats. The vast majority of AD experimental models almost exclusively consist of transgenic mice expressing human genes, resulting in amyloid plaques and NFT [66–68]. Other models have included invertebrate animals such as *Drosophila melanogaster* and vertebrates such as zebrafish. Considering the reduced relatedness in physiology between humans and these models, the latter is less extensively studied. [69,70].

Finding a naturally occurring AD model is appealing, as it would likely represent changes in human AD more accurately. Prior studies have demonstrated that gestational protein restriction leads to severe physiological and morphological changes in neurons [31,71–74] in addition to behavioral changes [75,76] and delays in cognitive and intellectual functions [77,78]. Thus, our results indicate that exposing the rat fetus to gestational protein restriction was sufficient to cause enhanced Aβ and tau protein hippocampal deposition in 88-week-old LP as compared to age-matched NP offspring.

Furthermore, changes in the cell numbers were observed, accompanied by hippocampal-dependent memory perturbations in LP progeny early than in control offspring. We suggest that these neural disorders may be key in causing the cognitive decline in early-stage AD-like rodent models.

Besides, consistent with previous studies, LP offspring significantly reduced lifespan (approximately 12 weeks) compared to NP offspring [79–81]. Incidentally, studies investigating the relationship among influence at the beginning of the developmental phase, fetal programming, and shortening of telomeres found each of these were strongly associated with perinatal stressors such as maternal stress, maternal food restriction, and maternal overnutrition [69–72].

Aging is characterized by a progressive loss of physiological integrity, leading to impaired function and increased vulnerability to death. Thus, comprehensive hallmarks of aging, including genomic instability, telomere attrition, epigenetic alterations, nutrient sensing, mitochondrial dysfunction, cellular senescence, stem cell exhaustion, and altered cellular communication, must be further investigated in this programmed experimental model. The current data suggest a high impact of maternal protein restriction on progeny lifespan and hippocampus structure, significantly accelerating the loss of hippocampal function, impacting learning/memory performance, and supposedly developing neurodegenerative-like syndromes in 88-week-old LP murine offspring. Carefully, we can suppose that maternal protein restriction in rats may be a probable, elegant, and novel method for constructing an AD-like model in adult male offspring.

## Acknowledgments and Financial support

This work was supported by Fundação de Amparo à Pesquisa do Estado de São Paulo (2013/12486-5), Coordenação de Aperfeiçoamento de Pessoal de Nível Superior (CAPES) and Conselho Nacional de Desenvolvimento Científico e Tecnológico (CNPq, 465699/2014-6).

## SUPPLEMENTAL INFORMATION

### METHODS

#### MWM proximity measure

The proximity measure optimized computer tracking to identify a rat’s position relative to the target location; it is considered highly sensitive to age-related learning/memory impairment in aged rats [24]. This study uses new proximity measures to the behavioral analysis of learning in the water maze task. We did this to test the memory/learning ability based on the proximity measure for use in characterizing individual differences in the effects of aging on spatial learning.

The need for such an index was based on the previously identified limitations of the customary analysis and on certain special features of the effects of aging on spatial learning ability.

The present study used the proximity of the animal’s position to provide several training trials and probe trial performance analyses. The proximity measure was obtained by sampling the animal’s position in the maze (10 times per second) to record its distance from the escape platform in 1-s averages.

By this method, scores obtained with the proximity measure are designed to reflect search error; they represent deviations from an optimal search, from a direct path to the goal (hidden platform) in the water maze.

The assessment of proximity to the target provides a more efficient analysis method than the multiple measures traditionally used to characterize probe trial performance: platform crossings and path length or time in the quadrant.

An analysis of young rat performance that used this measure showed rapid acquisition of improved search accuracy during the interpolated probe trials. Comparison of young and aged groups demonstrated that age was most pronounced relatively early in training because aged rats acquired a spatial bias more slowly. Thus, in this study, the probe trial analysis using proximity to the target was sensitive to an age-related impairment in spatial learning.

It appears that some rats become less proficient in learning the information that is required for efficient navigation to a specific location. So, the method used here offered a sensitive, efficient, and valid approach to assessing this age-related cognitive deficit.

## RESULTS (Supplemental information)

**Figure S1.**
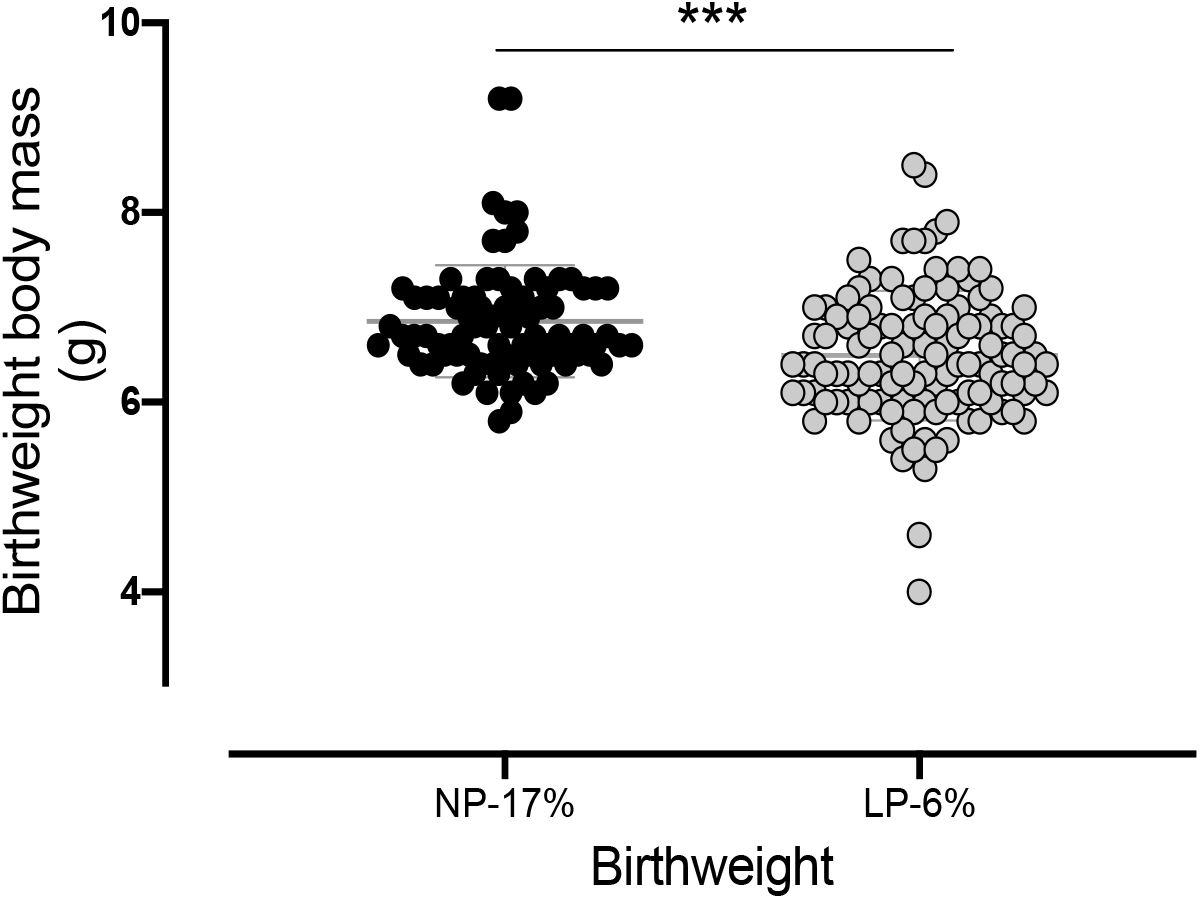
Offspring bodyweight (g) at birth NP progeny as compared to LP offspring. Results are depicted as scatter dot-plot and are expressed as means ± SEM; the comparisons involving only two means within or between groups were performed using a Student’s *t*-test. Welch’s *t*-test was used to correct situations characterized by heteroscedasticity. When statistically significant differences were indicated between selected means, posthoc comparisons were performed using Bonferroni’s contrast test. The level of significance was set at *p < 0.05.

**Figure S2.**
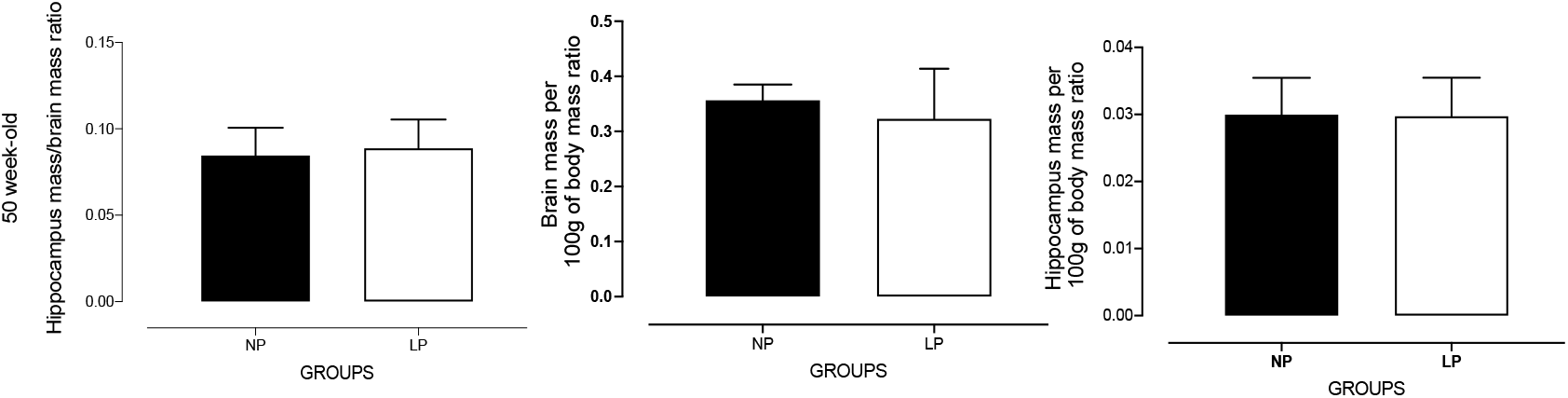
Offspring brain and hippocampal masses (g) in 50-week-old NP (n=5) compared to age-matched LP (n=5) offspring. Results are expressed as means ± SEM; comparisons involving only two means within or between groups were performed using a Student’s *t*-test. The level of significance was set at *p < 0.05.

